# Analytical Assessment of Metagenomic Workflows for Pathogen Detection with NIST RM 8376 and Two Sample Matrices

**DOI:** 10.1101/2024.09.24.614717

**Authors:** Jason G. Kralj, Stephanie L. Servetas, Samuel P. Forry, Monique E. Hunter, Jennifer N. Dootz, Scott A. Jackson

## Abstract

We assessed the analytical performance of metagenomic workflows using NIST Reference Material 8376 DNA from bacterial pathogens spiked into two simulated clinical samples: cerebral spinal fluid (CSF) and stool. Sequencing and taxonomic classification were used to generate signals for each sample and taxa of interest, and to estimate the limit of detection (LOD), the response function, and linear dynamic range. We found that the LODs for taxa spiked into CSF ranged from approximately (0.1 to 0.3) copy/μL, with a linearity of 0.96 to 0.99. For stool, the LODs ranged from (10 to 221) copy/μL, with a linearity of 0.99 to 1.01. Further, discriminating different *E. coli* strains proved to be workflow-dependent, as only one classifier:database combination of the three tested showed the ability to differentiate the two pathogenic and commensal strains. Surprisingly, when we compared the response functions of the same taxa in the two different sample types, we found those functions to be the same, despite large differences in LODs. This suggests that the “agnostic diagnostic” theory for metagenomics may apply to different target organisms *and* different sample types. Using RMs, we were able to generate quantitative analytical performance metrics for each workflow and sample set, enabling relatively rapid workflow screening before employing clinical samples. This makes these RMs a useful tool that will generate data needed to support translation of metagenomics into regulated use.

**Importance:** Assessing the analytical performance of metagenomic workflows, especially when developing clinical diagnostics, is foundational for ensuring that the measurements underlying a diagnosis are supported by rigorous characterization. To facilitate the translation of metagenomics into clinical practice, workflows must be tested using control samples designed to probe the analytical limitations (e.g. limit of detection). Spike-ins allow developers to generate fit-for-purpose control samples for initial workflow assessments and inform decisions about further development. However, clinical sample types include a wide range of compositions and concentrations, each presenting different detection challenges. In this work, we demonstrate how spike-ins elucidate workflow performance in two highly dissimilar sample types (stool and CSF); and we provide evidence that detection of individual organisms is unaffected by background sample composition, making detection sample agnostic within a workflow. These demonstrations and performance insights will facilitate translation of the technology to the clinic.

## Introduction

Next-generation sequencing-based metagenomics (a.k.a. metagenomic sequencing, MGS) pathogen detection offers tremendous promise as an “agnostic diagnostic,” enabling relatively unbiased sample characterization. (1, 2) This approach typically works by reducing samples to DNA/RNA, preparing them for a sequencer, collecting the raw sequence data for the sample, and then comparing the sequences to databases to assign likely identities of the sample constituents. This contrasts with *targeted* sequencing, which is effectively a highly multiplexed PCR assay that requires *a priori* knowledge of the targets. Data can be reanalyzed using multiple bioinformatics tools and/or databases, making comparisons and refinement of workflows possible without rerunning samples.

Clearly, different sample types can have dramatically different profiles, with a wide range of organism concentrations and diversity, (3-6) resulting in different biases to overcome. (Figure 1)

**Figure 1.**
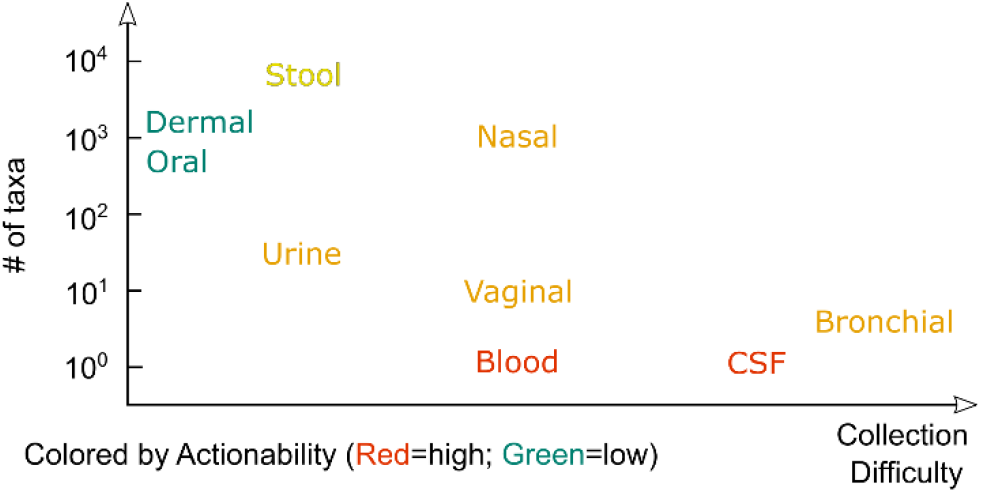
Human specimen typically investigated by MGS span a wide spectrum of complexity, difficulty of collection, and actionability. The number of taxa for a given sample type are approximate and may change as our understanding of each microbiome evolves. The actionability gives an approximate “score” for our understanding of the likelihood of finding pathogens at that site with potential for harm; individual use cases will vary.

For applications involving sterile fluids, the confirmed presence of any microbe would be cause for concern. In blood and CSF where samples should be nearly sterile, any DNA signal from a suspected pathogen above background is cause for concern, (3, 7-9) making LOD a critical parameter.

For samples with diverse microbial populations, only the presence of specific pathogenic strains would necessitate action. Such is the case in medium- to low-complexity samples (10s of taxa) such as urine or vaginal swabs. Here, having specificity for the suspected pathogen is a critical performance parameter, with LOD supporting the presence/absence, especially if attempting to identify antimicrobial resistant (AMR) strains. (4)

In high-complexity samples such as stool (100s to 1000s of species), (6) the analytical methods can dramatically impact the analysis. (10-13) Quantifying a critical abundance of specific strains such as shiga-toxin producers, viruses, & parasites, etc. may be higher priority than having a low LOD as with previous scenarios. Also, relative abundance of taxa may be clinically important. Therefore, basic analytical performance assessment, with some nuance specific for each application, remains a consistent need in all MGS applications.

Reference materials, used as spike-ins or internal controls, have begun to find utility to assess performance in metagenomics of MGS tools.(13, 14) In these studies, the response to spike-in concentrations in mock samples were assessed for 16S-based sequencing in performing absolute quantitation on samples. This marked a significant improvement to enable comparison studies and appears poised to enable reproducibility of metagenomics analyses.

These reference materials support workflow development by both facilitating the (a) assessment of analytical performance and (b) identification of analytical biases in the MGS workflow. With respect to analytical performance, the first steps include establishing the LOD, response linearity, and dynamic range; without establishing analytical performance characteristics, translation into a clinical setting becomes extraordinarily challenging. It is important to establish analytical performance for each MGS workflow due to the biases that result from each step (e.g. sample collection/handling, extraction, processing, and the bioinformatics); all of which result in a distortion of the original sample. Together, these biases cause a discrepancy between the reported and actual sample compositions, ultimately impacting workflow performance. Reference materials are useful in identifying these biases yet they have not yet been widely implemented. (7, 10, 12, 15, 16)

Individual laboratory assessments have previously shown that purpose-built workflows (i.e. fit-for-purpose for specific sample types) perform well on clinical sample types and have been approved for diagnostic use through CAP-CLIA. These laboratories have targeted clinical needs for detecting non-human DNA in otherwise sterile body fluids, specialize in given sample types, and have performed rigorous workflow characterization and validation. (3, 9, 17) However, to date, there are no examples of FDA-cleared testing devices, highlighting the difficulty in translating this diagnostic for broad use.

To aid translation of metagenomic workflows into regulated applications, there is clear need for methods assessing the analytical performance of a workflow using appropriate controls – organisms of interest, at appropriate levels, and in matrix. And though quantitative reference materials for MGS applications have been limited, the release of NIST RM 8376 aimed to remedy that. (18) This material can be employed to assess taxon-specific performance and identify potential weaknesses in a workflow before moving into clinical evaluation. (19)

In this work, we demonstrated how components of RM 8376 can be used to assess an MGS workflow’s limits of detection, linearity of response, and linear dynamic range in two different sample types, cerebral spinal fluid (CSF) and stool. The main goal of this publication was to demonstrate the utility of genomic reference materials and potential experimental designs; however, we also observed some interesting, and in some cases, unexpected results about common classifiers.

For CSF (low complexity and abundance), performance with both classifiers was the same. For stool (high complexity and abundance), one classifier had a significantly lower LOD for the variable spike-ins than the other; however, it failed to discriminate the strains of *E. coli*.

The discrimination (between *E. coli* strains) provided a greater challenge as compared to the first set of spike-ins due to subtle differences in the strains with potentially significant clinical implications. Ultimately, only 1 of the 3 workflows tested could achieve the required specificity.

Surprisingly, the response functions derived from the CSF and stool sample data suggest that MGS workflows, at least from library preparation through bioinformatics, were agnostic to the sample type, despite LODs differing by >100x in the two matrices. This would have significant implications for the utility of *in silico* experiments in early-stage studies and may be useful in *ex post facto* expanding the range of taxa from an approved workflow.

## Results & Discussion

### Experimental Overview

As part of its mission to promote U.S. innovation and industrial competitiveness, NIST has been working with stakeholders in industry, academia, and other government agencies to develop standards to support the advancement of MGS for clinical diagnostics and biosurveillance applications. As a result of this effort we developed RM 8376, a set of DNA from 19 different bacterial pathogens. (18) To demonstrate the utility of this material we present an experimental strategy for deploying these types of reference materials in the evaluation of an MGS workflow from library preparation through analytical performance assessment.

The strategy utilized RM 8376 as both internal controls (*A. hydrophila* and *L. pneumophila*) and variable spike-ins (*K. pneumoniae, E. faecalis, N. meningitidis*, and *L. monocytogenes*) in two clinically relevant matrices: CSF and stool. CSF typically contains low levels of human DNA and should not include significant levels of any microbial and/or viral nucleic acids. This makes it an ideal candidate for diagnostics based on metagenomics, as background signal is low, thereby simplifying the review of post-metagenomics analysis. Hence, CSF was selected as the low-complexity, low-background sample type for assessing RM 8376 spike-ins’ fitness for purpose for limit of detection and response-linearity characterization. In contrast, stool often contains a highly diverse population of microbes with an overall high density/abundance. Pathogenic strains of many enteric pathogens may differ only slightly in their genomes from otherwise commensal flora, making identification challenging. Additionally, these matrices represent extremes in terms of microbial abundance and complexity and served to bracket the range of clinical sample types described earlier (Figure 1).

Briefly, this experimental approach included addition of a subset of the strains of RM 8376 listed above into DNA extracted from pooled human CSF or stool. The internal controls were added to the extracted DNA, with individual spike-ins added in a log-fold dilution series to individual aliquots covering a 6-log range. With this setup, we could evaluate the analytical performance of the library preparation through bioinformatic analysis. Here, we focused on taxonomic classifiers and databases.

### Cerebral Spinal Fluid (CSF) Use Case

The determination of the LOD for each taxon consisted of 3 steps:

1. Estimate the background signal for each taxon and determine the minimum signal for detection/quantitation.
2. Perform a linear regression on the data above the minimum signal.
3. Use the linear model with the minimum signal to determine the LOD/LOQ.

We examined the background signal for all samples. (Figure 2, left) We observed the samples with high spike-in concentrations, which appeared to generate background normalized abundances much greater than low spike-in samples. We determined it was only appropriate to include samples with spike-in abundances less than 10^4^ copy/μL because these samples’ compositions were nearly the same and would more closely resemble a “typical” clinical sample (see SI). Typically, 0 or 1 absolute reads were observed for each taxon in any of these background samples, indicating the CSF samples were typically free of these species. From this, the minimum signal above background for each taxon was calculated. We note that the analytical performance metrics found in this use case apply only valid for each specific workflow. Significantly increasing the sequencing read depth 10- or even 100-fold would likely improve the LOD here because so few absolute reads were observed in the background; though, the per-sample cost may significantly increase as well.

**Figure 2.**
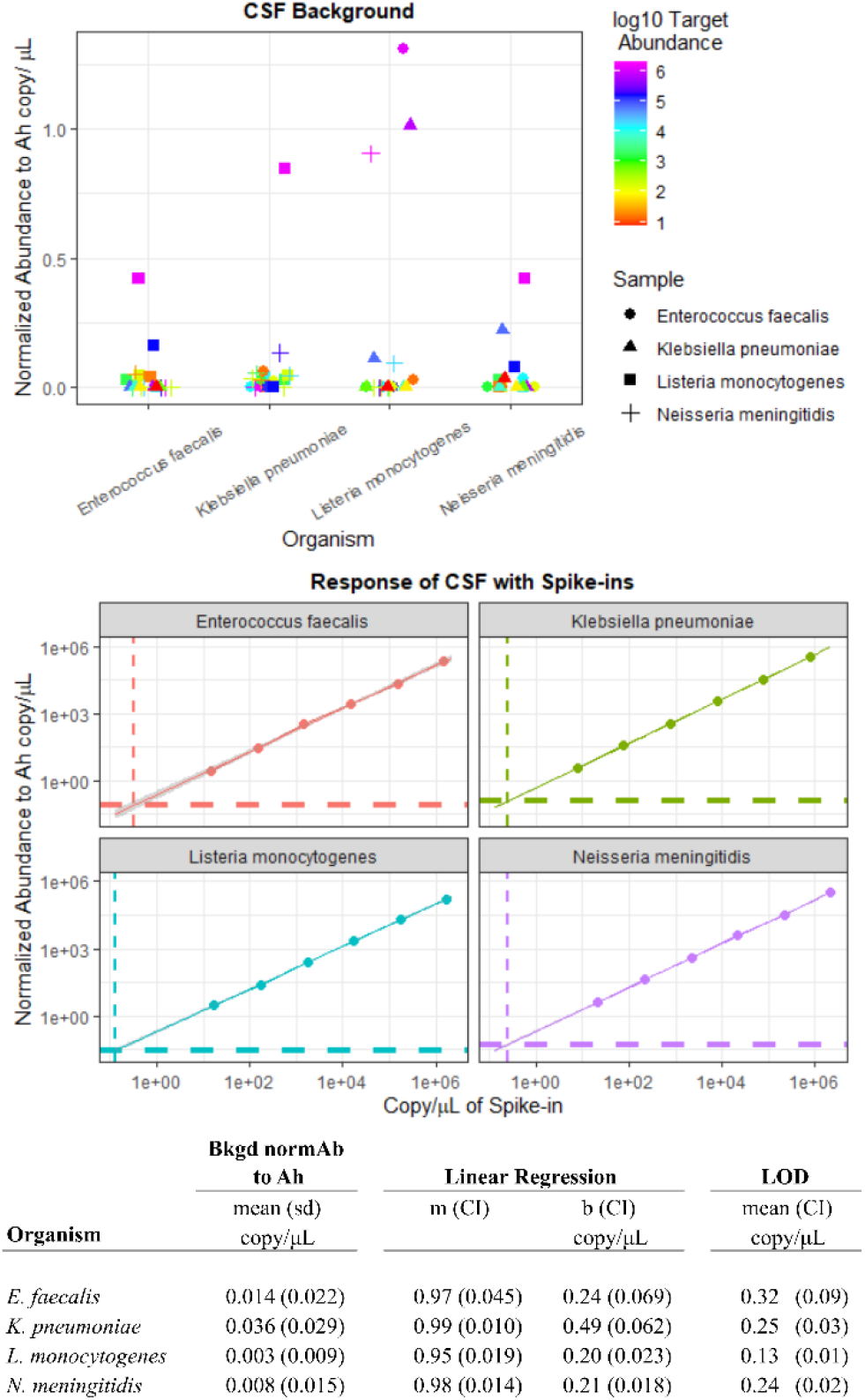
Individual organisms were spiked into CSF at different concentrations ranging from approximately (10^6^ to 10^1^) copy/μL. (Top) The background signal for each taxon was plotted as the normalized abundance to Aeromonas hydrophila (Ah). Control samples spiked with high amounts of other taxa (i.e. not the target taxa) generated significantly higher background signals compared with the control samples with low levels of spiked amounts. Because of this, we identified two criteria for including data in the LOD calculation – the target taxon could not be spiked into that sample, and the spiked-taxon concentration was below 10^4^ copy/μL (excluding blue and purple data points) so that the sample profile approximately resembles clinical samples with low levels of pathogens. (Middle) The response for each spiked-in taxon was plotted as normalized abundance to Ah vs. the spiked-in copy number. A linear regression with confidence band was included. The filtered background signal (horizontal dashed line) combined with the linear regression of the data were used to estimate the LOD for each organism (vertical dashed line). The LODs were similar for each organism, ranging from approximately (0.1 to 0.3) copy/μL. Subtle differences between the performance of each taxon were likely significant and due to how unique each read could be interpreted by the classifier, though this effect was small. (Bottom) Analytical performance metrics for the centrifuge classifier for CSF. The background signal relative to Ah, linear regression (b+m·log x), and LOD for the four taxa are shown here. The values are shown as mean (95 % confidence interval), unless otherwise noted. The slopes of the regression (m) showed a response of approximately 1 for the range of spike-in concentrations studied. The LODs correspond to sub-10-read signals, indicating that increasing the number of reads per sample would likely lower the LOD; however, this would require additional study, increase cost, and may not be necessary based on the particular criteria for a given use case.

A regression of the data above the minimum signal was then calculated and plotted. (Table 1, Figure 2 Right). For these 4 taxa, the slope of the response was approximately 1.0 with uncertainty of approximately 1 % to 5 % (see Figure 2, bottom), indicating no saturation of signal over the range of spike-in concentrations tested.

**Table 1.**
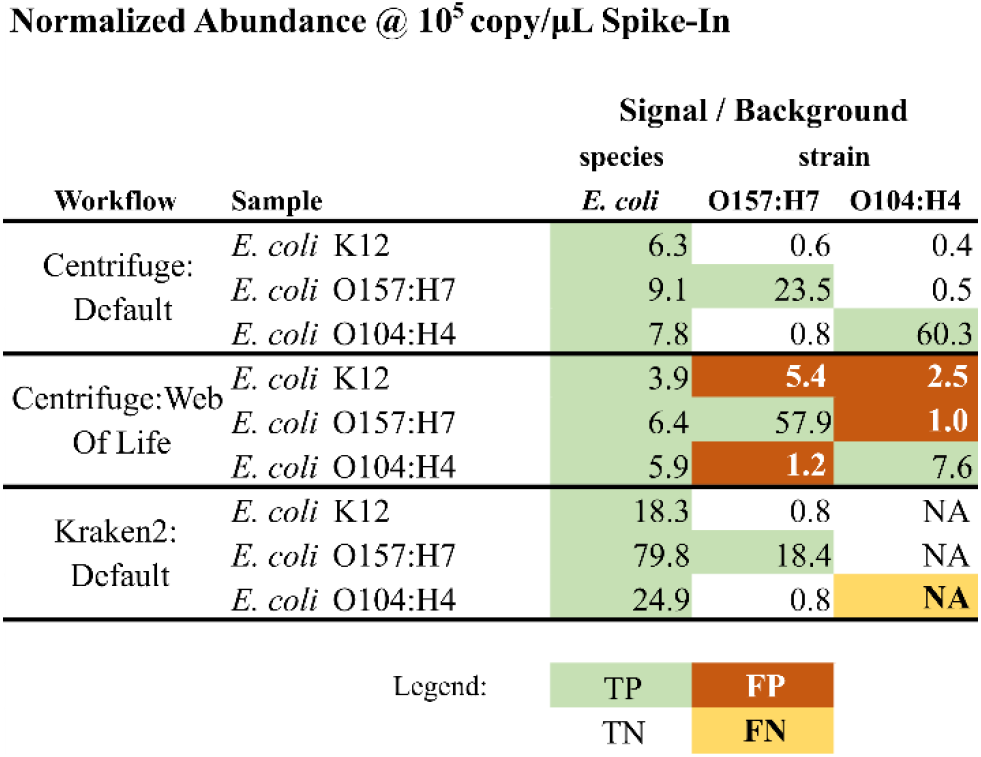
Analytical response of 3 E. coli strains spiked into stool at approximately 10^5^ copy/μL using the classifier centrifuge with two databases and kraken2. For ease of comparison, the Signal / Background is the measured normalized copy number concentration to the background concentration for each taxon. The default database correctly identified all 3 samples as having E. coli, and correctly discriminated the pathogenic strains from the commensal strain and each other. The WoL database also correctly identified all 3 samples as E. coli; however, the sample with the K12 strain was identified as having both O157:H7 and O104:H4 strains present. There was also positive signal crossover between pathogenic strain, though the off-target signal was signficantly lower than the on-target, which would be easily filtered. The kraken2:default workflow could not detect the E. coli O104:H4 strain, but could correctly discriminate the O157:H7 strain from the commensal and O104:H4 strains. The response curves for each sample and strain can be found in Fig.SI-3.

Normalized Abundance @ $10^5$ copy/ μL Spike-In
| Workflow | Sample | Signal / Background |  |  |
| --- | --- | --- | --- | --- |
|  |  | species | strain |  |
|  |  | *E. coli* | O157:H7 | O104:H4 |
| Centrifuge: Default | *E. coli* K12 | 6.3 | 0.6 | 0.4 |
|  | *E. coli* O157:H7 | 9.1 | 23.5 | 0.5 |
|  | *E. coli* O104:H4 | 7.8 | 0.8 | 60.3 |
| Centrifuge:Web Of Life | *E. coli* K12 | 3.9 | 5.4 | 2.5 |
|  | *E. coli* O157:H7 | 6.4 | 57.9 | 1.0 |
|  | *E. coli* O104:H4 | 5.9 | 1.2 | 7.6 |
| Kraken2: Default | *E. coli* K12 | 18.3 | 0.8 | NA |
|  | *E. coli* O157:H7 | 79.8 | 18.4 | NA |
|  | *E. coli* O104:H4 | 24.9 | 0.8 | NA |

Legend:
|  |  |
| --- | --- |
| TP | FP |
| TN | FN |

The linear model was used with the minimum signal to estimate the LOD for each taxon (Figure 2, bottom). The LODs ranged from approximately (0.1 to 0.3) copy/μL. These values were statistically different from each other, though of the same order of magnitude.

Linear dynamic range would typically also be assessed for where the response signal saturates or becomes unstable; however, we were unable to reach this maximum value. Since the typical operating range for this matrix is at or near the LOD, which is approximately 6 orders of magnitude below the maximum value we tested, we concluded that LDR was sufficient for low-complexity CSF samples.

The same data can be used to compare alternative workflows (i.e. quality filtering, subsampling, classifiers, databases) with respect to answering the question of which workflow is better suited for a given sample set. We assessed the performance using the classifier kraken2 (Figure SI-1) and compared the results to the workflow using centrifuge. The results for CSF data were nearly identical both in terms of LOD and linearity for the 4 taxa studied. Hence, either/both data workflows would be comparable in an investigation involving CSF with those taxa.

It is worth noting that this analysis accounts for performance of the sequencing and bioinformatics. Extraction efficiency is a known variable that impacts the response, and it may be necessary to increase the read depth of each sample to account for this in clinical samples. In this use case, each taxon appears to respond similarly. We would expect the response of most other bacterial DNA to be similar in CSF using the specific sequencer and experimental methods used. However, the first step here was to assess LOD and response using a quantified DNA standard--and to that end the experiment was successful in assessing these analytical performance metrics.

While we do not seek to establish this workflow as a diagnostic, evaluating the overall analytical performance in terms of comparability to existing workflows highlights the utility of reference materials for assessing comparability of workflows. And given the simplicity of using the quantified spike-ins, assessing pre-clinical analytical performance of the metagenomics portion of the workflow would inform on potential challenges (i.e. background signal, need for pre-amplification) before using patient samples.

One similar workflow from a previous clinical study estimated their LOD in CSF as between (0.1 to 0.2) genome copy/mL cell equivalents based upon 10 RPM-r (reads per million-relative to baseline), read length, and an average bacterial genome size of 4 Mbase.(9) However, two rounds of universal PCR were used to pre-amplify the targets, which we did not employ here, resulting in the 1000-fold lower detection threshold. Focusing on the 10 RPM-r LOD figure from the sequencing data alone, we achieved similar analytical sensitivity (approximately 3 RPM, based upon baseline estimate for minimum signal, LOD, and measured reads from samples) with this workflow. Hence, when accounting for the differences in sample preparation, the analytical performance derived from the contrived samples used here compared well with the clinical studies.

### Stool Use Case

The stool spike-in samples were analyzed using a similar workflow to the CSF samples. Briefly, we added two internal control organisms and variably spiked with a log-dilution of select RM 8376 components to extracted stool DNA. Those samples were sequenced and analyzed to look for the LOD and response of the spike-ins. We plotted the background signal and above-background response for the samples. (Figure 3)

**Figure 3.**
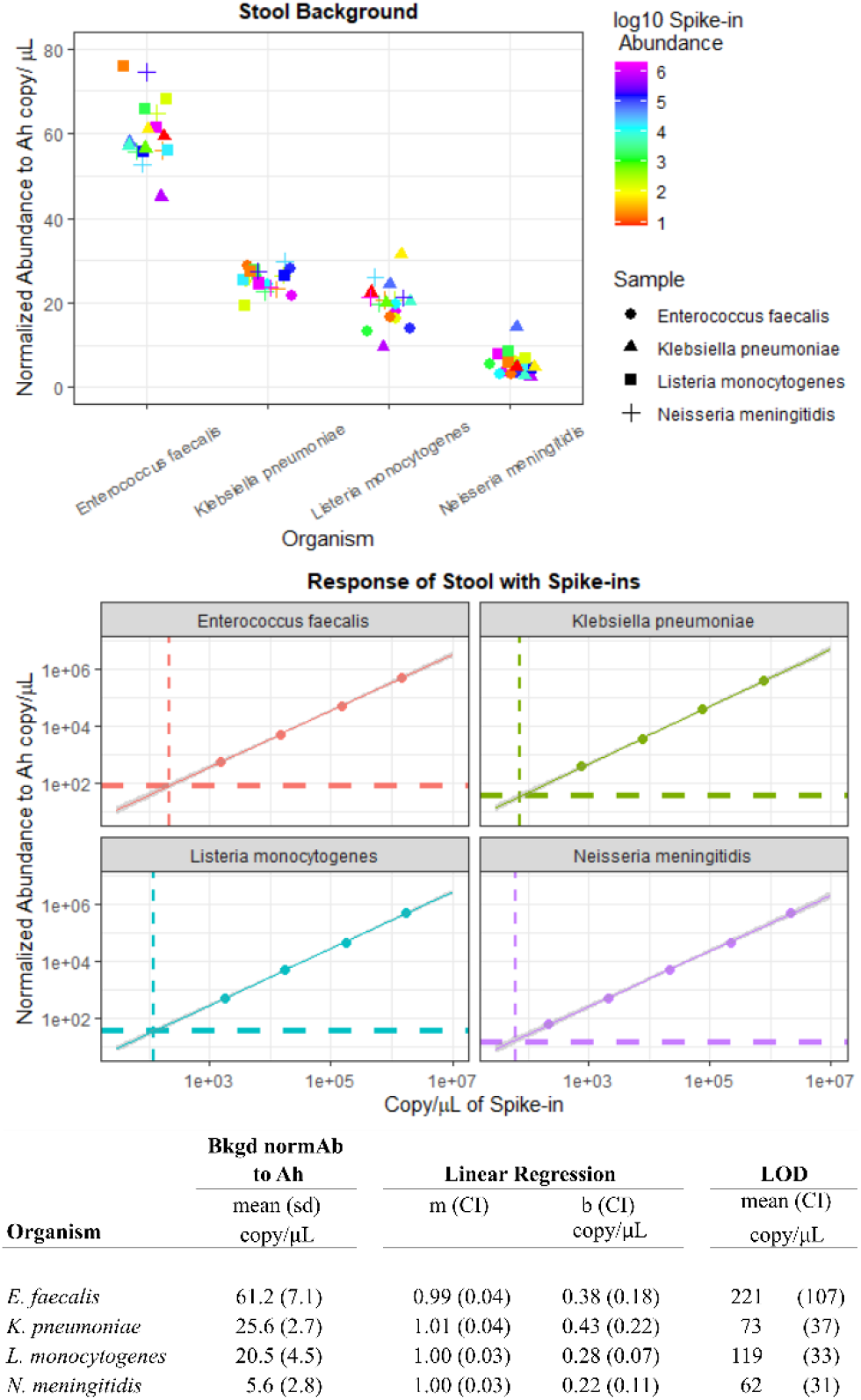
Individual organisms were spiked into human stool at different concentrations ranging from approximately (10^6^ to 10^1^) copy/μL. (Top) the background signal for each organism was assessed similar to the criteria used for CSF. The background signal for each taxon in stool samples was significantly higher than in CSF. This was expected due to the rich and diverse population of microbes found in stool, with some bias due to highly-abundance taxa (e.g. Enterococcus). Using a taxon-specific assessment, differences in performance were easily identified and this information could be used to properly establish LODs. (Middle) The response for each organism was plotted as the normalized abundance to Ah vs. the spike-in concentration as described in Figure 2. Here, both the magnitude and the differences in LOD by taxon were larger than with CSF, likely due to confounding signal from related taxa within the stool. (Bottom) Analytical performance metrics for the centrifuge classifier for stool. The same method was used as described in Figure 2. As expected, the background signal and LODs for each taxon in stool samples was significantly higher than for CSF. However, the linear regression for each organism had nearly identical parameters regardless of background, suggesting that for any given taxon, the workflow response is independent of the sample composition.

The background signal for the spike-in taxa was significantly higher in stool than for CSF, as will be shown later. We did not observe any significant change in background from high vs. low abundance spike-in samples; however, we only included samples with spike-ins less than 10^4^ copy/μL (n=12 per taxon) to be consistent for comparisons to CSF in later discussions. Also, unlike the CSF data, the spike-in concentrations span the LOD. The linear regression and LOD calculations were tabulated and compared with CSF (Figure 2, bottom).

The background data showed a clear difference in LOD for each taxon (Figure 3, top). We expected *Enterococcus* species could be present in the stool and generate significant background signal for that spike-in, and this was the case. The other 3 spike-ins also showed background signal. Given the broad diversity and high abundances of stool microbes, this result was not particularly surprising.

The kraken2 workflow results are shown in Figure SI-2. The response functions were similar between kraken2 and centrifuge. However, the LODs for the kraken2 workflow for these taxa were lower than centrifuge:default, indicating that alternative workflows/databases may improve performance. While kraken2 may outperform centrifuge for this sample set, both may be suitable for use if they meet the LOD needed for clinical utility; other factors including performance of a specific panel of organisms, speed, and ease of use may factor into the use of one/both/neither. We will explore this further in attempting to discriminate strains of *E. coli*.

### Demonstrating detection specificity of commensal vs. pathogen *E. coli*

Another test for MGS is to demonstrate the ability to discriminate organisms such as *C. difficile*, and various pathogenic strains of *E. coli*, etc. from commensal and/or non-pathogenic near-neighbor species or strains. Isolates can be sequenced deeply and assembled to paint a clear picture; however, this is time-consuming and expensive, and may not result in improved clinical outcomes. Instead, MGS-based systems that can relatively rapidly identify/discriminate pathogenic strain from whole stool directly, without isolation, would have clinical utility.

We tested the effect of using different databases with the same classifier and data, to see if the workflows would perform differently. For this demonstration, the classifier centrifuge was used with its default and the Web of Life (WoL) (20) databases, keeping all other parameters the same.

The goal of the analysis was to detect and differentiate *E. coli* sequences generated from the addition of commensal K12 strain DNA vs. two pathogenic strains in RM 8376 (an O157:H7 and O104:H4 strain). Since the workflow can report on both species- and strain-level detection, we set up our analysis to report on *E. coli* at the species level, and strains *E. coli* O157:H7 and *E. coli* O104:H4. All samples should contain *E. coli*, with only the pathogenic strains giving positive signals for their respective strain when present. We also chose to report the signal:background (similar to how others have normalized reporting (8)) because the background for *E. coli* in this sample was relatively high, making species- and strain-level normalized abundances appear disparate. (Table 1)

The results showed that the classifier Centrifuge:default correctly identified the addition of *E. coli* DNA above background in all samples, and only identified the correct pathogenic strains when present. In comparison, using the same workflow but with the WoL database did not result in discrimination of K12 *E. coli*. The K12 *E. coli* DNA gave a strong positive response for both pathogenic strains. Both pathogenic strains gave strong positive signals for their correct strain ID, with weak positive signals for the other. While the default database was able to discriminate pathogen vs non-pathogen, the WoL database failed the basic discrimination test.

In addition to testing a single classifier with two databases, we also evaluated performance of the kraken2 workflow with the default database. This workflow could not identify the O104:H4 strain despite its inclusion in the database, with the sample containing nearly 10^6^ genome copy/μL registering 0 reads. The workflow did show good discrimination of commensal and O157:H7 strains. However, this clear lack of response for O104:H4 (Figure SI-3) necessitates a different approach to meet the needs of this (*E. coli* discrimination) analysis. These data highlight the impact database selection can have on a workflow, and that taxonomic classifiers’ utility are strongly tied to this critical feature. Further, differences between kraken2 and centrifuge performance further highlights the need to examine the entire panel of target taxa when selecting a metagenomic workflow. While this does not represent an exhaustive list of potential pathogenic targets, it shows that the metagenomic workflow can discriminate these pathogenic and commensal strains. Hence, both the materials and the methodologies outlined here enabled a relatively simple and rapid assessment of fitness-for-purpose without employing any clinical specimens.

### Stool Analysis Final Thoughts

It is important to verify workflow performance for *each taxon* because it is critical to identify potential challenges before moving forward with clinical samples. The workflow response should not be extrapolated to other taxa (or other taxonomic classes). Though similar organisms may share similar performance, and performance from one organism may inform experimental design for another, the myriad factors previously discussed may improve or degrade detection and/or discrimination relative to other taxa.

## Effect of Different Sample Type on Analytical Performance of Individual Organisms

Examination of the linear models of each taxon revealed that the slopes and intercepts for individual taxa did not change between CSF and stool samples for the centrifuge:default (or kraken2:default, as shown in the SI) workflows (DNA library preparation through bioinformatics steps). This suggests that a given organism will produce the same analytical response when analyzed with a single MGS workflow regardless of sample matrix. Thus, the “agnostic diagnostic” theory for MGS-based systems may be extended from to the ability to identify any target within a sample to also include the analytical response of an organism across different sample types. However, we examined a limited sample set (i.e. CSF & stool), so additional work examining more sample types and complete workflows (i.e. including sample handling, DNA extraction) will be needed to test this hypothesis.

In contrast, we observed that the LODs for the spiked-in taxa were approximately 250- to 900-fold higher in stool than CSF with our workflow. This was expected, because measurement sensitivity is limited by the background signal, which varies by sample type. In this study, CSF presents a relatively low microbial biomass background, while stool contains a significantly larger (and more diverse) microbial population. Furthermore, specific organism LODs will be impacted by the variety of compositions found across clinical samples. Application-specific LODs should always be established experimentally for each organism.

## Conclusions

RM 8376 is a bacterial pathogen DNA based Reference Material that supports metagenomic analytic performance assessments. We demonstrated that RM 8376 can be spiked into a background of DNA from other clinically relevant samples to evaluate analytical performance of a metagenomic workflows from sequencing library preparation and sequencing through bioinformatics processing. Thus, this material is a critical component within the suite of reference materials needed to assess performance at different steps in the metagenomics workflow.

We demonstrated the utility of DNA spike-ins for rapid metagenomic analytical performance assessment in samples differing in complexity by generating quantitative LODs and response functions, and generating data that would lead to workflow optimization. For both low-diversity CSF samples and high-diversity stool samples, taxon-specific LODs were generated. As expected, the LODs for spike-in organisms were higher in stool than CSF. The analysis of CSF samples showed similar sub-1 genome copy/μL detection for all taxa, with similar analytical performance of the sequencing and informatics steps to previous work.

Within the framework of generating data supporting clinical utility, this approach generates an actionable pass/fail result for continued testing or redevelopment. If a workflow does not perform as expected on a DNA-only test set, it will likely perform worse when tested with whole-cell controls because additional variables (i.e. sample storage and nucleic acid extraction) negatively impact the workflow. Additional whole-cell reference materials and characterization would be required for those upstream steps, and a correction factor could be applied to the results generated here. Hence, DNA-based reference materials, such as RM 8376, support rapid analytical performance testing of metagenomic workflows.

RM 8376 includes several near-neighbor organisms to further assess metagenomic workflows for specificity. Different strains of *E. coli* were used and highlighted two key factors in workflow assessment. First, classifier and database selection impact analytical performance. Of the three workflows tested (kraken:default, centrifuge:default, and centrifuge:WebOfLife), only one could achieve the desired strain-level discrimination. Ideally, one would confirm the target taxa are contained within the database; however, even when this is the case (Web of Life), the classifier itself may not yield the desired performance/result. And second, each of the taxa, even down to the strain level, will have specific performance characteristics. The analytical performance results in this work should not be extrapolated to other taxa; though, the results may inform workflow developers and reviewers on the capabilities of a workflow and serve to identify extraordinary claims requiring additional investigation.

Metagenomic platforms hold great promise to fill a much-needed void for an “agnostic diagnostic.” Typically, this reference has referred to the system being target/pathogen agnostic; however, metagenomic workflows may go even further from an analysis standpoint, and also be sample agnostic—the signal/response of an individual organism may not depend on the background sample composition. Individual organism response functions for the CSF and Stool spike-in data showed remarkable similarity, despite organism LODs differing by several orders of magnitude. However, only two sample types were explored, evaluating only detection of DNA (i.e. not whole cells). We plan to study this further.

Looking ahead, tackling the challenge of transitioning metagenomics from a proof-of-concept laboratory test into regulated application requires careful step-by-step characterization and assessment of the entire metagenomic workflow. This requires a framework of validation studies, each utilizing reference materials that have been designed to assess performance covering each step in the workflow. In the work shown here, we highlighted the utility of DNA-based reference materials to characterize the analytical performance of taxonomic classifiers using two use cases. Building on the evidence found here, further studies employing *in silico* reference data could provide additional support for rapid workflow development and performance assessment. Reference data (i.e. genomes or raw sequencing reads) would provide the ground-truth for such experiments. Additionally, whole-cell reference materials would allow testing of additional experimental parameters including sample collection/storage and extraction. Together, this proposed suite of standards, combining reference materials & data, will help provide the experimental evidence for the translation of metagenomics into the regulated application space.

## Materials & Methods

### Cerebral Spinal Fluid (CSF)

The frozen CSF (BioIVT) obtained from 5 healthy volunteers was thawed on ice and extracted using the QIACube (Qiagen) with the Qiagen Blood and Tissue Kit (Cat #69504) and pooled. The CSF was separated into 12 180-μL aliquots, with elution of 100 μL each, pooled together totaling 1.2 mL at 0.023 ng/μL determined using the Denovix HS DNA kit (Wilmington, DE USA).

A 30 μL aliquot was removed for sequencing to serve as the CSF negative control (no spike-ins or internal controls). *A. hydrophila* and *L. pneumophila* from RM 8376 (NIST, Gaithersburg MD, USA) were added as internal controls to the mixture at 1E4 genome copy/μL and 1.4E3 genome copy/μL, respectively. 30 μL of this was removed for sequencing as the CSF with internal controls only.

Four components of RM 8376 were used as spike-ins to the stool DNA: *K. pneumoniae* (starting concentration 7.68E6 copy/μL), *N. meningitidis* (starting concentration 21.7E6 copy/μL), *E. faecalis* (starting concentration 14.8E6 copy/μL), and *L. monocytogenes* (starting concentration 17.4E6 copy/μL). Sample sets for each spike-in were generated using 1:10 serial dilutions (i.e. 5 μL *K. pneumoniae* stock into 45 μL CSF DNA, then 5 μL of that mix into 45 μL of CSF DNA, etc.) for 6 dilution sets of each spike-in. With 4 sample sets, two CSF controls, and a DNA elution buffer control, a total of 27 samples were then prepared for DNA shotgun sequencing.

### Stool

Two 1 mL vials (containing 100 mg/1 mL each) of homogenized candidate fecal reference material (BioIVT, Westbury NY, USA) were split into 4 aliquots each (250 µL) for extraction of genomic DNA using the Zymo *Quick*-DNA Fecal/Soil Microbe kit (KIT #D6012, Zymo Research) with 40 minutes bead-bashing at maximum speed (Vortex Genie, Scientific Industries, Inc., USA). The 8 DNA extractions were pooled and quantified at 7.0 ng/μL. The total volume was brought to 2.5 mL using 1× TE (Sigma-Aldrich, USA, Cat# 93302-500mL). Of this, 30 μL were withheld for sequencing as a negative control.

Two internal controls were added to the pooled DNA. Approximately 25 μL of *A. hydrophila* DNA and 2.5 μL of *L. pneumophila* DNA to achieve 10^5^ genome copy/μL and 1.4.10^4^ genome copy/μL, respectively. Another 30 μL of this was withheld for sequencing as stool DNA with internal controls.

Four components of RM 8376 were used as spike-ins to the stool DNA: *K. pneumoniae* (starting concentration 7.68E6 copy/μL), *N. meningitidis* (starting concentration 21.7E6 copy/μL), *E. faecalis* (starting concentration 14.8E6 copy/μL), and *L. monocytogenes* (starting concentration 17.4E6 copy/μL). Sample sets for each spike-in were generated using 1:10 serial dilutions (i.e. 5 μL *K. pneumoniae* stock into 45 μL stool DNA, then 5 μL of that mix into 45 μL of stool DNA, etc.) for 6 dilution sets of each spike-in.

For the *E. coli* experiments, two components of RM 8376 were used as spike-ins to the stool DNA: *E. coli* O157:H7 (starting concentration 8.84E6 copy/μL) and *E. coli* O104:H4 (starting concentration 8.86E6 copy/μL). Sample sets for each spike-in were generated using 1:10 dilutions as previously described for 6 dilution sets of each spike-in. The *E. coli* K12 DNA was obtained from Sigma-Aldrich (MBD0013, approximately 10 ng/μL, estimated starting concentration 1.7e6 copy/μL). The 5-sample set consisted of first diluting the stock 1:20, then 4 additional 1:10 dilutions.

### Sequencing protocols

Samples were prepared using the New England Biolabs (Ipswich MA, USA) NEBNext Ultra II FS kit (NEB #E7805, E6177) and the NEBNext Multiplex Oligos for Illumina (Dual Index Primers Set 1) (E7600S) following the manufacturer’s instructions. For CSF samples, the DNA concentration was measured using the Denovix dsDNA HS for each sample after library preparation and found to range between 34 ng/μL to 81 ng/μL for the spiked-in samples, 17.5 ng/μL for the CSF negative control, 80 ng/μL for the CSF with internal controls, and 0.9 ng/μL for the elution buffer. For stool samples, each sample was analyzed for DNA length and concentration using the Tapestation D5000 (Agilent, Santa Clara, CA), and pooled equally. For sequencing, the sample pool was diluted to 12 pmol/L.

The DNA were sequenced using an Illumina MiSeq (San Diego, CA) with a MiSeq Reagent Kit v2 (300 cycle) as 2×150 paired reads (Cat #MS-102-2002) following the manufacturer’s protocol, including PhiX controls at 12.5 pmol/L.

## Data Availability and Analysis

### Raw Data

The raw, dehosted (human) sequencing data can be found at the NCBI SRA under PRJNA1074773.

### Bash scripts

There were three bash scripts used to analyze the fastq files (SI-Bash Scripts). In Script #1, samples were de-hosted using bowtie2 against the human genome assembly GRCh38. In Script #2, we performed quality filtering (bbduk), taxonomic classification using centrifuge with the default database,(21) centrifuge with the Web of Life database,(20) and kraken2 with the default database.(22) Centrifuge outputs were converted to a kraken-style output using centrifuge-kreport. (SI-Bash Scripts)

The third bash script queried the kreports for reads of the specific taxa of interest using a list of organisms of interest and storing this as a <sample>.csv file.

The <sample>.csv files were then used as inputs in an R-script to perform final normalized abundance calculations and plotting.

### Normalizing Abundance

Normalized abundance was selected over relative abundance, reads per million (RPM), and reads per kilobase per million reads (RPKM). Because the normalization factor (Ah concentration) was known, the response from each taxon of interest can be quantified *independently* of other taxa.(15, 23) This would not be the case with the other metrics, where bias from every organism impacts the reported metric.

We calculated the normalized abundance for each taxon in each sample relative to *Aeromonas hydrophila* (Ah). The internal control concentration of Ah in CSF (10^4^ copy/μL) and stool (10^5^ copy/μL) was held constant for each sample type. The formula to calculate normalized abundance was:

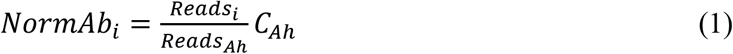

Where *NormAb*_*i*_ is the normalized abundance of species i, *Reads*_*i*_ were the assigned reads by the taxonomic classifier to species i, and *C*_*Ah*_ was the concentration of Ah.

In the experimental design, we also included *Legionella pneumophila* (Lp) as a second internal control. The Ah/Lp ratio (approximately 10) could be used to check for contaminating Ah and/or Lp that would impact normalization (>10 indicating background Ah, and <10 indicating background Lp). In this study, Ah and Lp background levels were not significant compared to the internal control abundance.

## Post-sequencing Analysis

### Minimum Signal

For each taxon, the background normalized abundance was estimated using samples where (1) the taxon was not spiked-in, and (2) only samples with spike-in abundances below 10^4^ copy/μL. For each taxon, this resulted in approximately 12 samples. This ensured that the sample compositions were similar and not overwhelmed by spike-in reads.

The background normalized abundance mean and standard deviation (sd) were calculated for each taxon. We set the minimum signal for finding the LOD (mean + 3 sd).

### Linearity

For each taxon, we filtered data for spike-ins with normalized abundance above the minimum signal. We fit the log-transformed data to a linear model log (NormAb) = m log (spike-in) + b. The 95 % confidence interval on the slope and intercept were estimated using a Student t-test.

### Limit of Detection (LOD)

For each taxon, we established the LOD by solving the linear model at the minimum signal for the spike-in concentration. The LOD confidence interval was estimated using the expanded uncertainty with the coefficient of variabilities of the slope and intercept of the model.

## Disclaimer

Certain equipment, instruments, software, or materials are identified in this paper in order to specify the experimental procedure adequately. Such identification is not intended to imply recommendation or endorsement of any product or service by NIST, nor is it intended to imply that the materials or equipment identified are necessarily the best available for the purpose.

## References

1. Gauthier NPG, Chorlton SD, Krajden M, Manges AR. 2023.. Agnostic Sequencing for Detection of Viral Pathogens. Clin Microbiol Rev 36:e0011922.

2. Govender KN, Street TL, Sanderson ND, Eyre DW. 2021.. Metagenomic Sequencing as a Pathogen-Agnostic Clinical Diagnostic Tool for Infectious Diseases: a Systematic Review and Meta-analysis of Diagnostic Test Accuracy Studies. J Clin Microbiol 59:e0291620.

3. Blauwkamp TA, Thair S, Rosen MJ, Blair L, Lindner MS, Vilfan ID, Kawli T, Christians FC, Venkatasubrahmanyam S, Wall GD, Cheung A, Rogers ZN, Meshulam-Simon G, Huijse L, Balakrishnan S, Quinn JV, Hollemon D, Hong DK, Vaughn ML, Kertesz M, Bercovici S, Wilber JC, Yang S. 2019.. Analytical and clinical validation of a microbial cell-free DNA sequencing test for infectious disease. Nat Microbiol 4:663–674.

4. Bloom SM, Mafunda NA, Woolston BM, Hayward MR, Frempong JF, Abai AB, Xu J, Mitchell AJ, Westergaard X, Hussain FA, Xulu N, Dong M, Dong KL, Gumbi T, Ceasar FX, Rice JK, Choksi N, Ismail N, Ndung’u T, Ghebremichael MS, Relman DA, Balskus EP, Mitchell CM, Kwon DS. 2022.. Cysteine dependence of Lactobacillus iners is a potential therapeutic target for vaginal microbiota modulation. Nat Microbiol 7:434–450.

5. Eisenstein M. 2020.. The skin microbiome. Nature 588:S209.

6. Arumugam M, Raes J, Pelletier E, Le Paslier D, Yamada T, Mende DR, Fernandes GR, Tap J, Bruls T, Batto J-M, Bertalan M, Borruel N, Casellas F, Fernandez L, Gautier L, Hansen T, Hattori M, Hayashi T, Kleerebezem M, Kurokawa K, Leclerc M, Levenez F, Manichanh C, Nielsen HB, Nielsen T, Pons N, Poulain J, Qin J, Sicheritz-Ponten T, Tims S, Torrents D, Ugarte E, Zoetendal EG, Wang J, Guarner F, Pedersen O, de Vos WM, Brunak S, Doré J, Weissenbach J, Ehrlich SD, Bork P. 2011.. Enterotypes of the human gut microbiome. Nature 473:174–180.

7. Kumar MSEV; Okrah, K; Hicks SC; Hannenhalli S; Cordava Bravo, H. 2018.. Analysis and correction of compositional bias in sparse sequencing count data. BMC Genomics 19:799.

8. Miller S, Naccache SN, Samayoa E, Messacar K, Arevalo S, Federman S, Stryke D, Pham E, Fung B, Bolosky WJ, Ingebrigtsen D, Lorizio W, Paff SM, Leake JA, Pesano R, DeBiasi R, Dominguez S, Chiu CY. 2019.. Laboratory validation of a clinical metagenomic sequencing assay for pathogen detection in cerebrospinal fluid. Genome Res 29:831–842.

9. Wilson MR, Sample HA, Zorn KC, Arevalo S, Yu G, Neuhaus J, Federman S, Stryke D, Briggs B, Langelier C, Berger A, Douglas V, Josephson SA, Chow FC, Fulton BD, DeRisi JL, Gelfand JM, Naccache SN, Bender J, Dien Bard J, Murkey J, Carlson M, Vespa PM, Vijayan T, Allyn PR, Campeau S, Humphries RM, Klausner JD, Ganzon CD, Memar F, Ocampo NA, Zimmermann LL, Cohen SH, Polage CR, DeBiasi RL, Haller B, Dallas R, Maron G, Hayden R, Messacar K, Dominguez SR, Miller S, Chiu CY. 2019.. Clinical Metagenomic Sequencing for Diagnosis of Meningitis and Encephalitis. N Engl J Med 380:2327–2340.

10. Ji Y, Huotari T, Roslin T, Schmidt NM, Wang J, Yu DW, Ovaskainen O. 2020.. SPIKEPIPE: A metagenomic pipeline for the accurate quantification of eukaryotic species occurrences and intraspecific abundance change using DNA barcodes or mitogenomes. Mol Ecol Resour 20:256–267.

11. Knudsen BE, Bergmark L, Munk P, Lukjancenko O, Prieme A, Aarestrup FM, Pamp SJ. 2016.. Impact of Sample Type and DNA Isolation Procedure on Genomic Inference of Microbiome Composition. mSystems 1.

12. Li M, Tyx RE, Rivera AJ, Zhao N, Satten GA. 2022. What Can We Learn about the Bias of Microbiome Studies from Analyzing Data from Mock Communities? Genes (Basel) 13.

13. Tourlousse DM YS, Ohashi A, Matsukura S, Noda N, Sekiguchi Y. 2017.. Synthetic spike-in standards for high-throughput 16S rRNA gene amplicon sequencing. Nucleic Acids Res 45:e23.

14. Venkataraman A, Parlov M, Hu P, Schnell D, Wei X, Tiesman JP. 2018.. Spike-in genomic DNA for validating performance of metagenomics workflows. BioTechniques 65:315–321.

15. McLaren MR, Willis AD, Callahan BJ. 2019.. Consistent and correctable bias in metagenomic sequencing experiments. Elife 8.

16. Morton JT, Marotz C, Washburne A, Silverman J, Zaramela LS, Edlund A, Zengler K, Knight R. 2019.. Establishing microbial composition measurement standards with reference frames. Nat Commun 10:2719.

17. Gu W, Deng X, Lee M, Sucu YD, Arevalo S, Stryke D, Federman S, Gopez A, Reyes K, Zorn K, Sample H, Yu G, Ishpuniani G, Briggs B, Chow ED, Berger A, Wilson MR, Wang C, Hsu E, Miller S, DeRisi JL, Chiu CY. 2021.. Rapid pathogen detection by metagenomic next-generation sequencing of infected body fluids. Nat Med 27:115–124.

18. Kralj J, Tourlousse D, Hunter M, Romsos E, Toman B, Vallone P, Jackson S. 2022.. Reference Material 8376 Microbial Pathogen DNA Standards for Detection and Identification. Commerce Do, NIST Publications, https://www.nist.gov/publication.

19. Kralj JG, Servetas SL, Forry SP, Jackson SA. 2020.. Considerations for performance metrics of metagenomic next generation sequencing analyses. BioRxiv doi:10.1101/2020.12.17.423212.

20. Zhu Q, Mai U, Pfeiffer W, Janssen S, Asnicar F, Sanders JG, Belda-Ferre P, Al-Ghalith GA, Kopylova E, McDonald D, Kosciolek T, Yin JB, Huang S, Salam N, Jiao JY, Wu Z, Xu ZZ, Cantrell K, Yang Y, Sayyari E, Rabiee M, Morton JT, Podell S, Knights D, Li WJ, Huttenhower C, Segata N, Smarr L, Mirarab S, Knight R. 2019.. Phylogenomics of 10,575 genomes reveals evolutionary proximity between domains Bacteria and Archaea. Nat Commun 10:5477.

21. Kim D, Song L, Breitwieser FP, Salzberg SL. 2016.. Centrifuge: rapid and sensitive classification of metagenomic sequences. Genome Res 26:1721–1729.

22. Wood DE, Lu J, Langmead B. 2019.. Improved metagenomic analysis with Kraken 2. Genome Biol 20:257.

23. Zaramela LS, Tjuanta M, Moyne O, Neal M, Zengler K. 2022.. synDNA-a Synthetic DNA Spike-in Method for Absolute Quantification of Shotgun Metagenomic Sequencing. mSystems doi:10.1128/msystems.00447-22:e0044722.

