## Supplemental Information for "Analytical Assessment of Metagenomic Workflows for Pathogen Detection with NIST RM 8376 and Two Sample Matrices"

**Selection of background samples to match "typical" clinical samples**

The pooled CSF samples sequence data were composed almost entirely human reads, even after subtracting them. This is expected for healthy CSF, which should be a sterile fluid containing only host DNA. In cases of individuals with infected CSF, NGS-based detection would still likely reveal the vast majority of reads to be human. Hence, even infected (clinically-positive) samples will almost always contain relatively low bacterial DNA burden.

We designated negative control samples for a taxon as any sample where that taxon was not spiked in, but may contain other taxa. For example, a CSF spiked with *Klebsiella* would count as a negative control for *Neisseria*. However, we limited the selection of negative controls to those samples with relatively low spike-in abundance to more accurately mimic clinical samples. This reduced the potential number of negative controls, but provided a stricter (and we feel, more appropriate) criteria for inclusion.

In addition, we observed that the absolute number of reads in negative control samples for each taxon were low (typically 0 or 1). We chose to exclude any samples with high spike-in abundance from being a negative control because they did not represent a typical clinical sample profile. High spike-in samples contained mostly spike-in reads, diluting the internal control Ah (by approximately 100- to 1000-fold) and effectively amplifying any background signal. In a clinical device, there would likely be other strategies put in place to deal with this, including setting a minimum number of reads for the total sample, internal control(s), and/or taxa of interest.

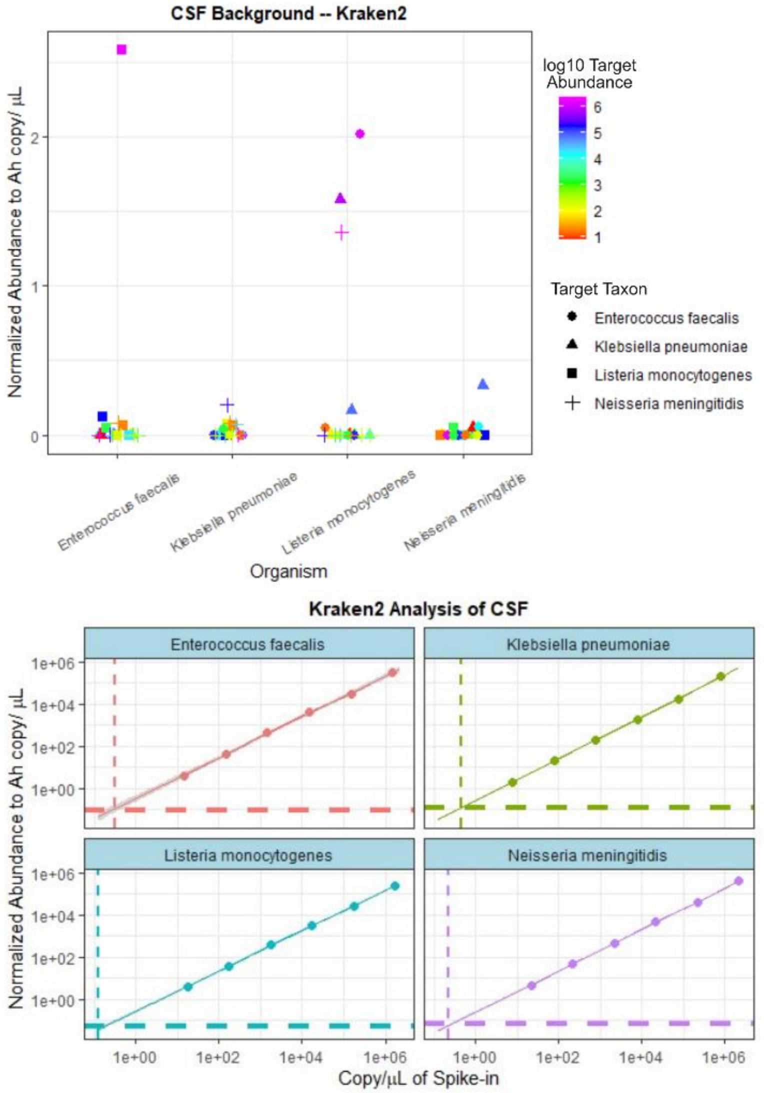


| Organism | Bkgd normAb to Ah<br>mean (sd)<br>copy/µL | Linear Regression<br>m (CI) | <br>b (CI)<br>copy/µL | LOD<br>mean (CI)<br>copy/µL |
|---|---|---|---|---|
| *E. faecalis* | 0.014 (0.028) | 0.97 (0.039) | 0.24 (0.110) | 0.32 (0.09) |
| *K. pneumoniae* | 0.025 (0.032) | 0.99 (0.011) | 0.26 (0.021) | 0.44 (0.03) |
| *L. monocytogenes* | 0.006 (0.017) | 0.95 (0.016) | 0.20 (0.032) | 0.13 (0.02) |
| *N. meningitidis* | 0.010 (0.021) | 0.98 (0.013) | 0.21 (0.022) | 0.23 (0.02) |

*Figure SI-1 Kraken2 workflow analysis of the CSF data -- individual organisms were spiked into CSF at different concentrations ranging from approximately $10^6$ to $10^1$ copy/µL. (Top) The background signal for each taxon was plotted as the normalized abundance to Aeromonas hydrophila (Ah), as shown in previous figures.*

*(Middle) The response for each spiked-in taxon was plotted as normalized abundance to Ah vs. the spiked-in copy number as shown in previous figures.*

*(Bottom) Analytical performance metrics for the kraken2 classifier for CSF, as described previously. The linear regression and LODs are nearly identical to the values calculated for the centrifuge workflow.*

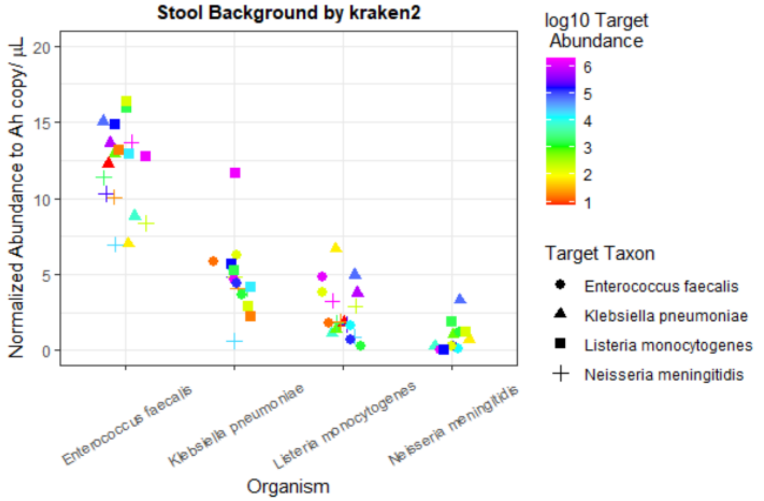


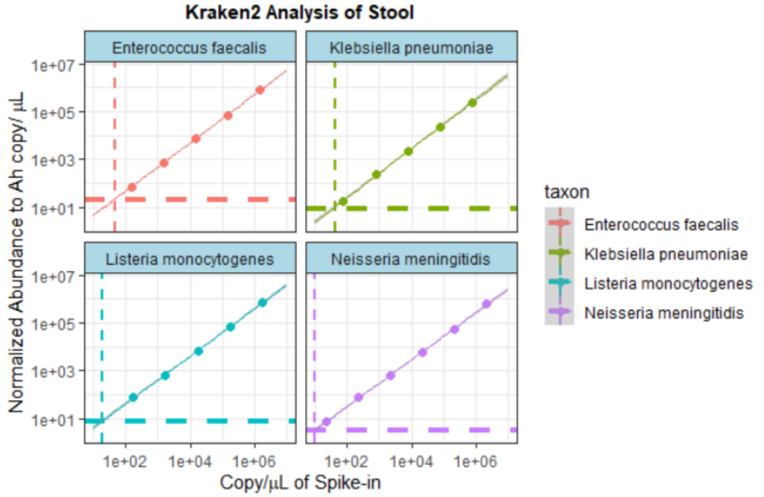


| Organism | Bkgd normAb mean (sd) copy/µL | | Linear Regression m (CI) | Linear Regression b (CI) copy/µL | LOD mean (CI) copy/µL |
|---|---|---|---|---|---|
| *E. faecalis* | 11.7 | (3.1) | 1.01 (0.02) | 0.46 (0.13) | 44 (12) |
| *K. pneumoniae* | 4.2 | (1.4) | 1.02 (0.05) | 0.23 (0.11) | 34 (17) |
| *L. monocytogenes* | 2.4 | (1.8) | 0.99 (0.03) | 0.43 (0.13) | 18 (5) |
| *N. meningitidis* | 0.6 | (0.9) | 0.98 (0.04) | 0.34 (0.14) | 10 (4) |

*Figure SI-2 Kraken2 workflow on stool samples – (top) background signal for each taxa shows the differences in response of each organism, as described previously.*

*(middle) The response for each spiked-in taxon was plotted as normalized abundance to Ah vs. the spiked-in copy number. The same method was used as described previously.*

*(Bottom) Assessing the background signals and response of the 4 spike-ins using kraken2. The LODs of the target taxa using kraken2 were significantly lower than using centrifuge.*

*Table SI-1 Assessing the stool background signals and response of the 4 spike-ins using two taxonomic classifier kraken2. The same conventions were used as described in Figure SI-1 (bottom). The LODs of the target taxa were significantly lower than using centrifuge. However, other factors may impact selection, including the performance of specific organisms, ease of use, etc.*

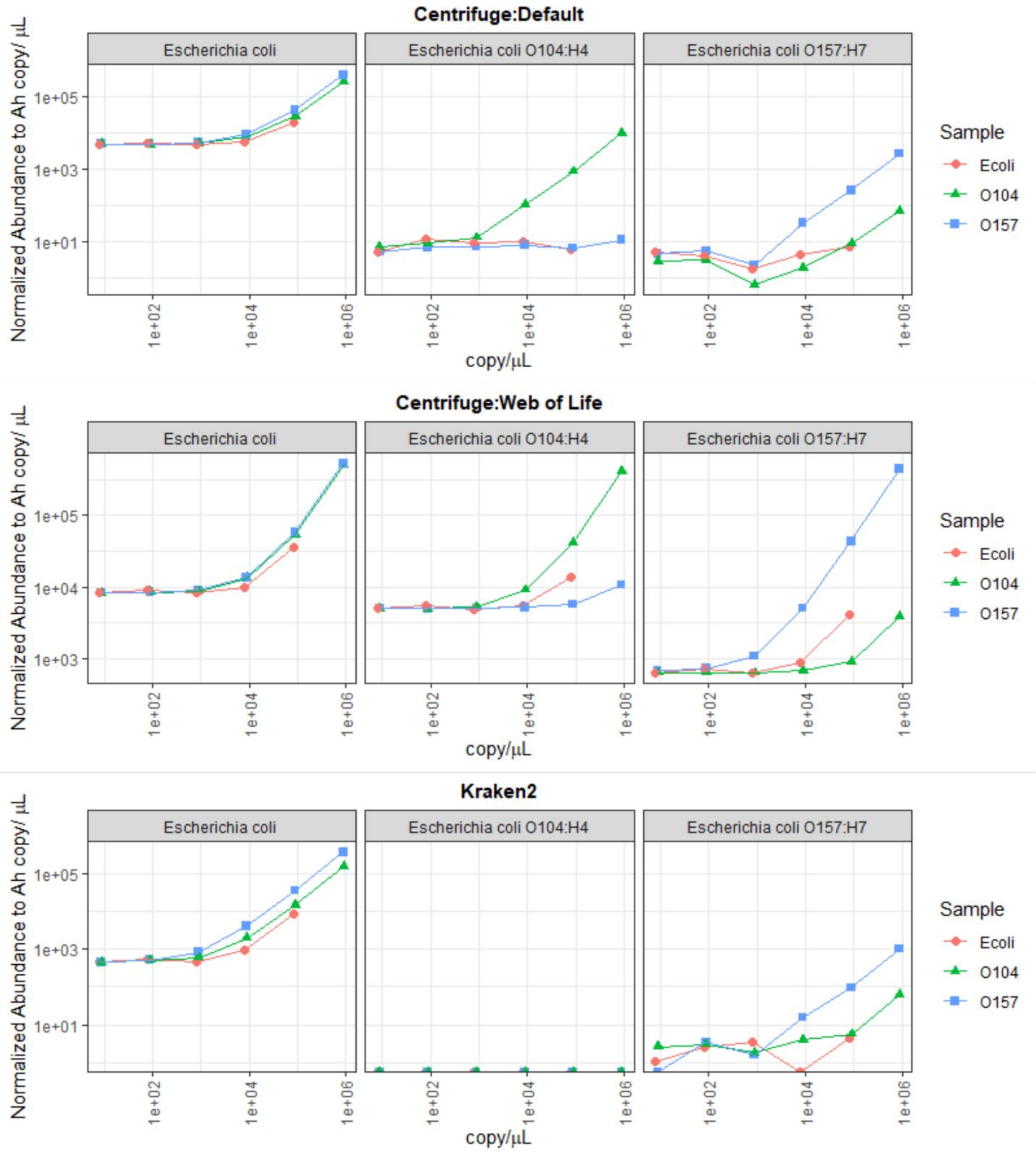


*Figure SI-3 The response curves for all 3 classifier:database combinations. Centrifuge:default shows the expected behavior – discrimination of commensal E. coli from the two pathogenic strains. Centrifuge:WoL shows that the commensal E. coli gives signal in both pathogenic strains, incorrectly indicating it is pathogenic. Kraken2 shows that commensal E. coli does not appear as a pathogenic strain; however, the O104:H4 strain is not detected despite being included in the database, and can even appear incorrectly as O157:H7 at high abundance.*

### Bash Scripts

#### Script #1 -- Dehosting

```
#!/bin/bash

eval "$(conda shell.bash hook)"
   conda activate bowtie2-env

# Location of reference genome
host="/path/GRCh38_noalt_as/GRCh38_noalt_as"

cd raw_reads

while read samples; do

echo -e "\033[92mProcessing ${samples} \033[37m"
bowtie2 -p 36 -x $host \
        -1 ${samples}_L001_R1_001.fastq.gz \
        -2 ${samples}_L001_R2_001.fastq.gz \
        --un-conc-gz \
        ${samples}_host_removed > /dev/null

#       --met-file ${samples}_met.txt \

#       Standard cleanup
mv ${samples}_host_removed.1  ./host_removed/${samples}_host_removed_R1.fastq.gz
mv ${samples}_host_removed.2  ./host_removed/${samples}_host_removed_R2.fastq.gz

done <../samples

cd -
conda deactivate
```

#### Script #2 – Sample processing

```
#!/bin/bash

# VARIABLES & FILES MANAGEMENT #

# Subsampling #
  minlength=100

# File names #
  RAWREADS="raw_reads"
  OUTPUT="processedReads"
  subsampled="$OUTPUT/subsampled"
  samples="samples"

# Directories #
  mkdir -p ${OUTPUT}/subsampled
  mkdir -p results/{centrifugeWoL,centrifuge,kraken2}

# /FILES #
main_ReadProcessing(){

sampling
trimming
#subsampling
centrifuge_wol
centrifuge_default
kraken2_default
}

sampling(){

cd ${RAWREADS}
```

```
   ls *R1*.fastq.gz | sed s/_L001_R1_001.fastq.gz// > ../samples

cd -
}

trimming(){
# you will need to generate a text file called "samples" with the sample names.
# the switches you can adjust as necessary, but usable as is.

#files=$(ls -d $RAWREADS/*R1*.fastq.gz | sed s/_L001_R1_001.fastq.gz// | awk '{print $NF}')
cd $RAWREADS

echo "Processed Reads for " $PWD > processedReads.txt

while read sampleID; do
echo "Trimming ${x}..."
/path/bbmap/38.90/bbmap/bbduk.sh -Xmx84g \
  in1=${sampleID}_L001_R1_001.fastq.gz \
  in2=${sampleID}_L001_R2_001.fastq.gz \
  out1=../$OUTPUT/${sampleID}_R1.fastq \
  out2=../$OUTPUT/${sampleID}_R2.fastq \
  t=30 \
  qtrim=rl \
  trimq=20 \
  minlen=$minlength \
  maxns=0 \
  rieb=t |& tee -a ${sampleID}_bbduk_rpt.txt

# Generate processed reads report
  echo -e -n ${sampleID} '\t' | tee >> processedReads.txt
  sampleReads=$(grep "^@M" ../$OUTPUT/${sampleID}_R?.fastq | wc -l)
  echo $sampleReads | tee >>processedReads.txt


done < ../$samples

#  >> processedReads/${x}_bbduk.log 2>&1

# copy this info into the results files for R
cp processedReads.txt ../results/centrifuge/
cp processedReads.txt ../results/centrifugeWoL/
cp processedReads.txt ../results/kraken2/
cd -
}

subsampling(){

cd $OUTPUT

   N=500000     #### Set the number of subsampled reads ####

while read y;
 do
echo "Subsampling ${y}..."
        seqtk sample -s100 ${y}_R1.fastq $N > $subsampled/${y}_500K_R1.fastq
        seqtk sample -s100 ${y}_R2.fastq $N > $subsampled/${y}_500K_R2.fastq
        cat $subsampled/${y}_500K_R1.fastq $subsampled/${y}_500K_R2.fastq > $subsampled/${y}_combined_1M.fastq

done <../$samples
cd -

}

centrifuge_wol(){

databaseWoL="/path/WoLr1"
# add line to generate report in folder for database, numreads, etc.
echo "Centrifuge:WoL"

while read sampleID; do
```

```
echo "${sampleID}..."

centrifuge \
-x $databaseWoL \
-1 $OUTPUT/${sampleID}_R1.fastq \
-2 $OUTPUT/${sampleID}_R2.fastq \
-p 36 \
-S /dev/null \
--report-file ./results/centrifugeWoL/${sampleID}_centrifuge.txt

REPORT="./results/centrifugeWoL"

# Sort the output from centrifuge using the numReads
  (head -n 1 $REPORT/${sampleID}_centrifuge.txt && \
   tail -n +2 $REPORT/${sampleID}_centrifuge.txt | sort -nr -k5 -t$'\t') \
   > $REPORT/${sampleID}_centrifuge_sorted.txt


done < $samples
}

centrifuge_default(){

databaseDefault="/path/p+h+v"
#add report in folder for database and numreads

while read sampleID; do

echo "Centrifuge:default ${sampleID}..."

### centrifuge, run subsampled reads

centrifuge \
-x $databaseDefault \
-1 $OUTPUT/${sampleID}_R1.fastq \
-2 $OUTPUT/${sampleID}_R2.fastq \
-p 36 \
-S /dev/null \
--report-file results/centrifuge/${sampleID}_centrifuge.txt

REPORT="results/centrifuge"

# Sort the output from centrifuge using the numReads
(head -n 1 $REPORT/${sampleID}_centrifuge.txt && \
 tail -n +2 $REPORT/${sampleID}_centrifuge.txt | \
 sort -nr -k5 -t$'\t')  > $REPORT/${sampleID}_centrifuge_sorted.txt

done < $samples

}

kraken2_default(){

db_default="/path/kraken2-default"

export PATH="//builds/kraken2/scripts:$PATH"
pwd
while read x; do

echo "kraken2 processing ${x}..."

# confidence changed 0.001-->0.05 on 2021-10-29; should result in lower sens
#  but higher spec. of low-abundance taxa

kraken2 --db $db_default \
 --threads 36 \
 --use-names \
 --confidence 0.05 \
 --report ./results/kraken2/${x}_k2_report.txt \
```

```
 --paired \
 $OUTPUT/${x}_R1.fastq \
 $OUTPUT/${x}_R2.fastq \
 --output ./results/kraken2/${x}.txt

cut -f1-3,6-8 ./results/kraken2/${x}_k2_report.txt > ./results/kraken2/${x}_stdK2_report.txt
done < $samples
}

### Call the main list to execute ###
main_ReadProcessing
```

#### Script #3 – query for specific taxa

```
#!/bin/bash
# When calling script, you must specify the normalizing taxa (e.g. Aeromonas),
#               otherwise it will default to Homo sapiens.
# List of taxa from elsewhere or just by nano > organisms.

norm=${1:-"Homo sapiens"}
organisms="organisms"
# Default normalization to "Homo sapiens"
echo `pwd` "norm = ${1}" > abundance_list.csv

while read sample; do

normcount=$(awk -F '\t' -v NAME="$norm" '$4=="S" && $6 ~ NAME  {count+=$2} END {print count}' ${sample}_k2_report.txt)
# echo "$1, $normcount" >> abundance_list.csv

while read taxa; do
 awk -F '\t' -v SAMPLE="$sample" -v NAME="$taxa" -v nc="$normcount" \
 '$4=="S" && $6 ~ NAME  {count+=$2} END {print SAMPLE","NAME","count","count/nc}' \
 ${sample}_k2_report.txt | tee -a  abundance_list.csv

done <$organisms
done <../../samples
```
